# Fitting mechanistic epidemic models to data: a comparison of simple Markov chain Monte Carlo approaches

**DOI:** 10.1101/110767

**Authors:** Michael Li, Jonathan Dushoff, Ben Bolker

**Keywords:** MCMC, HMC, TSIR, Dispersion, Moment-matching

## Abstract

**Background:** Simple mechanistic epidemic models are widely used for forecasting and parameter estimation of infectious diseases based on noisy case reporting data. Despite the widespread application of models to emerging infectious diseases, we know little about the comparative performance of standard computational-statistical frameworks in these contexts. Here we build a simple stochastic, discrete-time, discrete-state epidemic model with both process and observation error and use it to characterize the effectiveness of different flavours of Bayesian Markov chain Monte Carlo (MCMC) techniques. We use fits to simulated data, where parameters (and future behaviour) are known to explore the limitations of different platforms and quantify parameter estimation accuracy, forecasting accuracy, and computational efficiency across combinations of modeling decisions (e.g. discrete vs. continuous latent states, levels of stochasticity) and computational platforms (JAGS, NIMBLE, Stan).

**Results:** Models incorporating at least one source of population-level variation (i.e., dispersion in either the transmission process or the observation process) provide reasonably good forecasts and parameter estimates, while models that incorporate only individual-level variation can lead to inaccurate (or overconfident) results. Models using continuous approximations to the transmission process showed improved computational efficiency without loss of accuracy.

**Conclusion:** Simple models of disease transmission and observation can be fitted reliably to simple simulations, as long as population-level variation is taken into account. Continuous approximations can improve computational efficiency using more advanced MCMC techniques.

## 1 Introduction

Simple homogeneous population models have been widely used to study emerging infectious disease outbreaks. Although such models can provide important insights — including estimated epidemic sizes and predicted effects of intervention strategies, as well as short-term forecasts — they neglect important spatial, individual-level and other heterogeneities. Decades of work have created frameworks that enable researchers to construct models that capture many of these more realistic aspects of infectious disease epidemics. But many challenges remain. In particular, estimating parameters (and associated uncertainties) is always challenging, especially for models incorporating multiple forms of heterogeneity, and especially during the early stages of an epidemic when data are limited. Using complex models that are insufficiently supported by data can lead to imprecise and unstable parameter estimates (Ludwig and Walters, 1985) — in such cases, researchers often revert to simpler models for practical purposes.

In the past few decades, researchers have begun to adopt Bayesian approaches to disease modeling problems. Bayesian Markov Chain Monte Carlo (MCMC) is a powerful, widely used sampling-based estimation approach. Despite the widespread use of MCMC in epidemic modeling (Morton and Finkenstädt, 2005; O’Neill, 2002), however, there have been relatively few systematic studies of the comparative performance of statistical frameworks for disease modeling O’Neill et al. (2000).

In this paper, we apply relatively simple MCMC approaches to data from simulated epidemics that incorporate stochasticity in both transmission and observation, as well as variable generation-interval distributions (not assumed to be known when fitting). We compare model approaches of varying complexity, including an estimation model that matches the simulation model. For each model we quantify parameter estimation accuracy and forecasting accuracy; this sheds light on which phenomena are most important to include in models to be used for estimation and forecasting.

We also compare three different MCMC platforms: JAGS (Plummer et al., 2003), NIMBLE (de Valpine et al., 2016) and Stan (Carpenter et al., 2016). In principle, for any given model, any valid method of MCMC sampling should eventually converge on the same (correct) posterior distribution. However, even with the relatively simple models considered here, a theoretically valid software package can experience problems in practice: we wanted to investigate this phenomenon. Furthermore, even when different platforms converge to essentially the same result, they may show large differences in computational efficiency: we therefore also quantify efficiency for the models we study.

## 2 Methods

We generated test data using a simple framework that combines a *transmission process* based on a simple discrete-time model with an *observation process* to account for incomplete reporting. Both processes are assumed to be stochastic. We then fit the observed cases from these simulations using Bayesian methods that model the underlying true number of infections as a latent (i.e., unobserved) variable. Our Bayesian fitting models explore an approach that matches the assumptions of the simulation model, as well as various simplifications: in particular, we explore simpler methods of accounting for variation in both the transmission process and the observation process, and the use of continuous rather than discrete latent variables. For simplicity, we have here assumed that data are reported on the same discrete time scale on which the disease process is simulated (but not that the reported period is the same as the generation time of the disease; see below). This assumption requires that the generation time be at least as the reporting period. It would be relatively straightforward to relax this assumption, for example by assuming that the epidemic dynamics occur on a finer time scale than the reporting interval, or by simulating in continuous time but fitting with a discrete-time model; we do not explore these questions here.

### 2.1 Simulation Model

The transmission process of our dual-process framework is based on the Reed-Frost chain binomial model, which can also be described as a discrete-time, stochastic compartmental SIR model (Ludwig, 1973). To account for the possibility that some fraction of the population may be beyond the scope of the epidemic — geographically or socially isolated, genetically resistant, vaccinated or immune due to previous exposure — we assume that only a proportion *P*_eff_ of the total census population is actually susceptible to infection. We further assume that, in every time step, only a proportion (randomly chosen with mean *P*_rep_) of new infections are actually observed. We model both transmission and observation using a beta-binomial (rather than binomial) distribution to account for additional sources of variation (i.e., overdispersion) in both processes. The equations are:
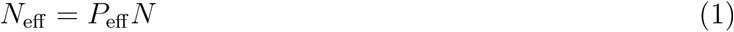

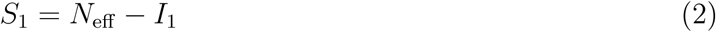

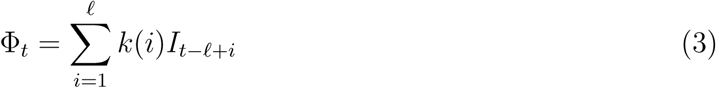

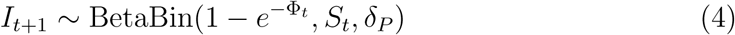

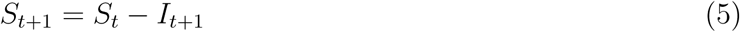

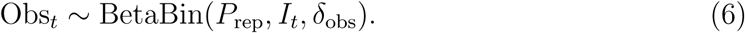
where Φ*_t_* is the force of infection at time *t*; *N*_eff_ is the effective population size; and ℓ is the number of lags.

The most common parameterization of the beta-binomial comprises three parameters: the binomial size parameter *N* plus two additional shape parameters (*α* and *β*) that describe the Beta distribution of the per-trial probability. Uses of the beta-binomial in statistical modeling instead typically transform the shape parameters into a pair of parameters that describe the per-trial probability and a dispersion parameter Morris (1997); larger values of the dispersion parameter *δ* correspond to less variability. We use a slight modification of this parameterization (see figure 1)

**Figure 1:**
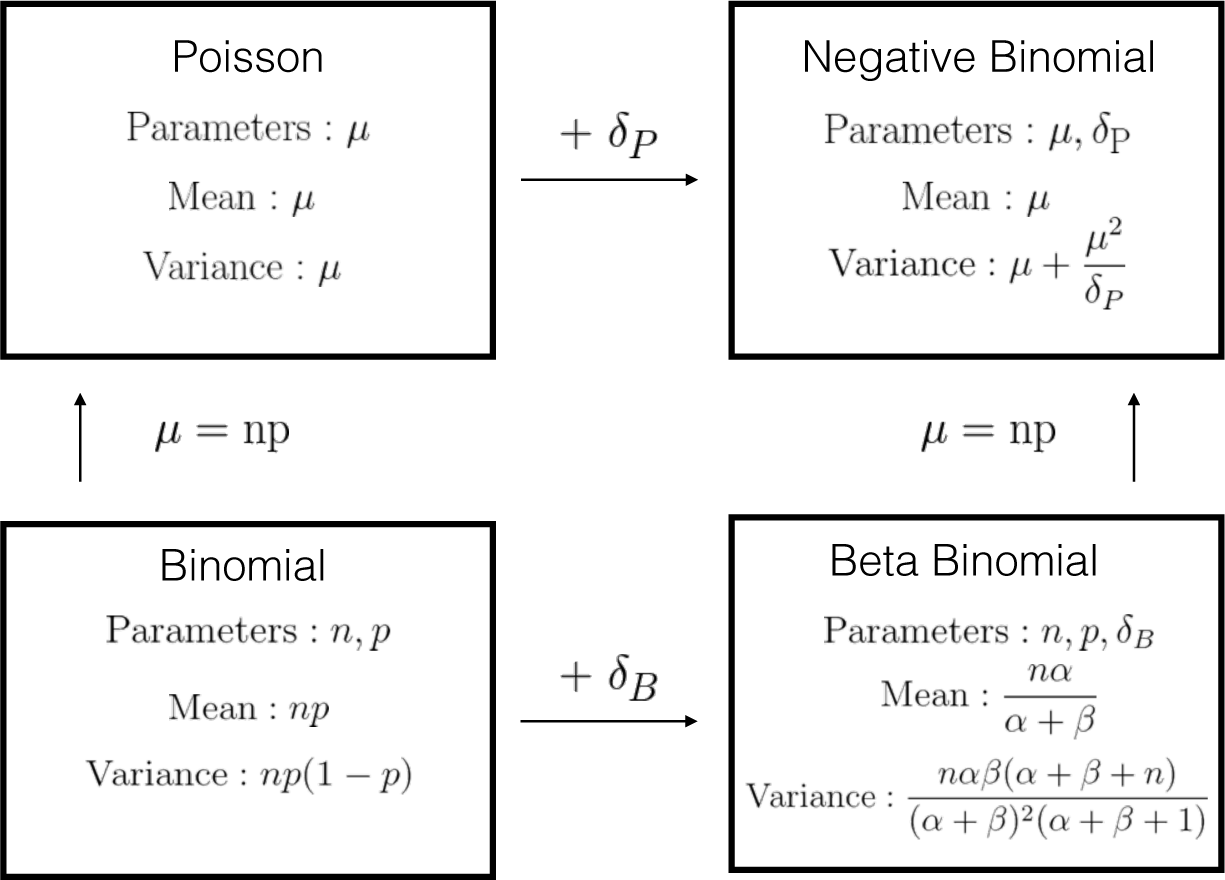
Discrete distribution relationships. For beta-binomial distribution (bottom right panel), we used an alternative parameterization *α* and *β*, where 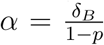 and 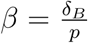. Moving from the top to bottom row adds a size parameter (replacing *μ* with *np*). Moving from left to right adds a dispersion parameter *δ_P_* and *δ_B_* for Poisson and Binomial distribution respectively.

We extend the Reed-Frost model by allowing the infectious period to last longer than one step, and the infectivity to vary based on how long an individual has been infected; we do this by parameterizing a transmission kernel that describes the force of infection coming from individuals who were infected ℓ time steps ago. For convenience, we assumed a fixed maximum window length (ℓ = 5). We then based our transmission kernel on a negative binomial distribution, truncated to fit this window:
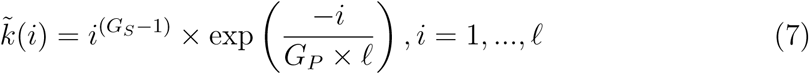

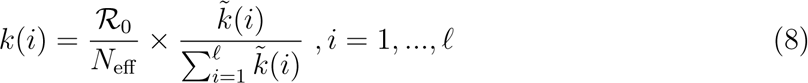

Here, 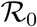 represents the basic reproductive number and *G_S_* and *G_P_* are shape and position parameters, respectively.

### 2.2 Fitting Model

#### 2.2.1 Transmission and Observational Process Errors

The transmission (eq. 4) and observation (eq. 6) processes in the simulation model are both defined as beta-binomial (BB) processes. In fitting, we used the BB to match the simulation model, but also tried several simpler alternatives: binomial (B), Poisson (P), and negative-binomial (NB) processes. Process B does not allow for overdispersion, while NB does not incorporate the size of the pool from which a value is chosen; that is, it is theoretically possible for a NB sample of the number of infections to be larger than the current susceptible population (although this is extremely unlikely when the *per capita* infection probability is small). Process P neglects both of these phenomena. Figure 1 illustrates the relationship of the four discrete distributions.

#### 2.2.2 Multiple Scale Decorrelation

The proportion of the population assumed to be effectively susceptible (*P*_eff_) and the reporting proportion (*P*_rep_) have very similar effects on observed incidence. We therefore reparameterized the model so that it uses a single parameter *P*_effrep_ for their product, and a second to govern how the product is apportioned between the two quantities:
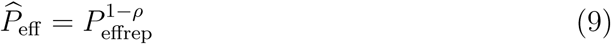

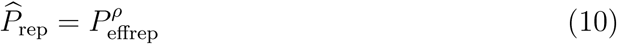

We expected *a priori* that this parameterization would improve statistical convergence, since it makes it possible to sample different values of the poorly constrained value of *ρ* without changing *P*_effrep_. It is straightforward to back-calculate *P*_eff_ and *P*_rep_ once the model is fitted. For similar reasons, we experimented with measuring infected individuals on a “reporting” scale in our continuous-variable models (see below).

#### 2.2.3 Continuous latent variables

Another simplification we considered was treating the unobserved number of underlying cases as a continuous variable. To do this, we matched the first two moments of the discrete distribution to a Gamma distribution (Figure 2).

**Figure 2:**
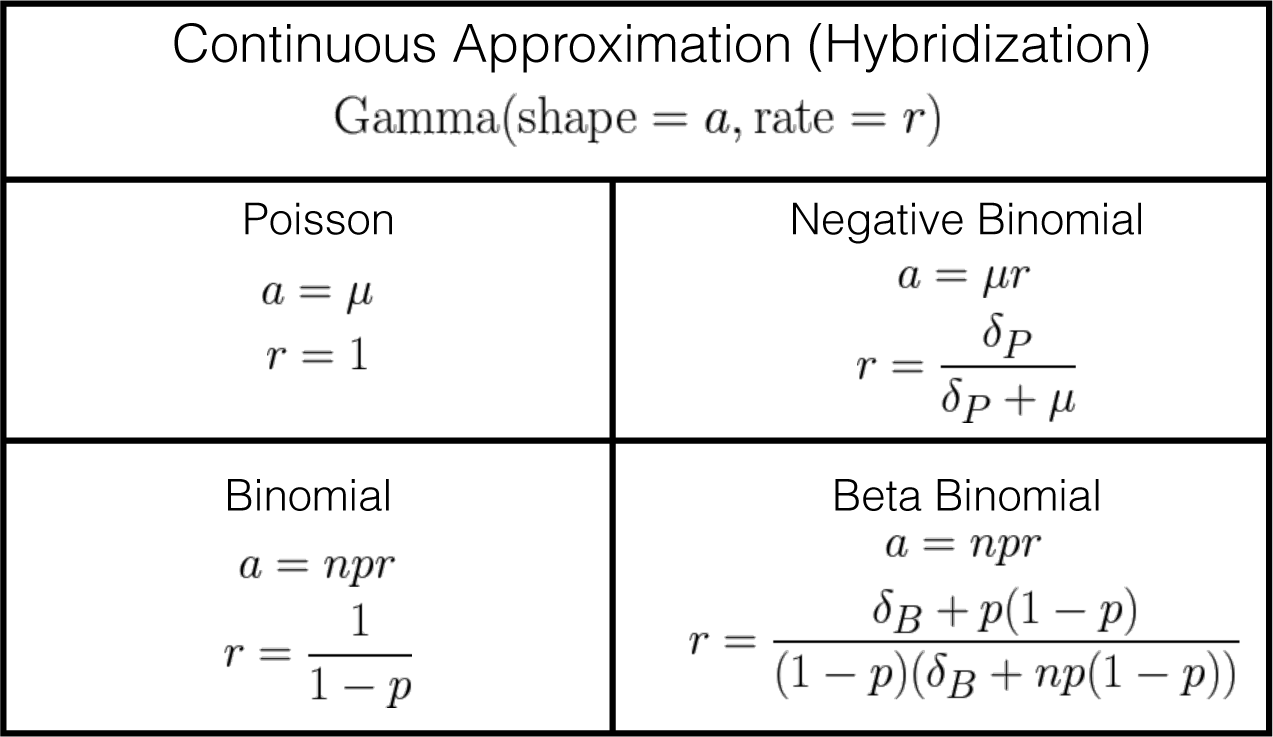
Continuous approximation of discrete distributions via moment matching. Distributions in Figure 1 were matched to a Gamma distribution with equivalent first and second moments.

Eq. 4 and 6 can be rewritten as:
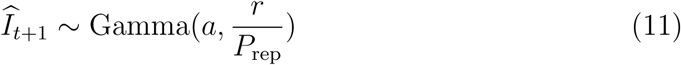

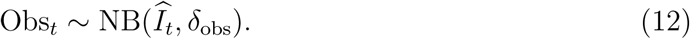

One advantage of this continuous approximation approach is that it allows us to scale our latent variable to help with model convergence, so that infected individuals are measured on the reporting scale. Another advantage is that it allows us to use MCMC sampling procedures such as Hamiltonian Monte Carlo (HMC), which cannot easily use discrete latent variables.

### 2.3 Bayesian Markov Chain Monte Carlo

In Bayesian MCMC, model parameters are sampled from the posterior distribution by a reversible Markov chain whose stationary distribution is the target posterior distribution. Classical MCMC techniques include the Metropolis-Hasting algorithm (Hastings, 1970), Gibbs sampling (Geman and Geman, 1984), and slice sampling (Neal, 2003). Recently, convenient implementations of a powerful MCMC technique called Hamiltonian Monte Carlo (HMC: also called hybrid MC) (Duane et al., 1987) have become available. HMC uses the concept of Hamiltonian dynamics to create a proposal distribution for the M-H algorithm, together with the leap-frog algorithm and the No U-Turn sampler (Hoffman and Gelman, 2014). HMC requires more computational effort per sample step compared to other MCMC techniques, but because subsequent steps are less strongly correlated it also produces more effective samples per sample step (Carpenter et al., 2016; Hoffman and Gelman, 2014).

#### 2.3.1 Platforms

Many software platforms implement the automatic construction of MCMC samplers for user-defined models. One of the most widely used platforms is JAGS (Just a Gibbs Sampler); despite its name, it combines a variety of MCMC techniques to fit models. NIMBLE (Numerical Inference for Statistical Models for Bayesian and Likelihood Estimation) is a more recent platform that allows users to flexibly model and customize different algorithms and sample techniques for MCMC. Neither JAGS nor NIMBLE has yet implemented HMC. One of the relatively few platforms that currently implements HMC is Stan, which provides full Bayesian inference for continuous-variable models based on the No-U-Turn sampler, an adaptive form of HMC.

#### 2.3.2 Simulation and Evaluations

A typical frequentist statistical simulation scheme fits multiple realizations to data generated from a fixed set of parameters that is determined *a priori* and evaluates the match of the parameter estimates to the true values. When we fit our Bayesian model using informative priors, frequentist coverage is generally higher than nominal values (i.e. 90% posterior intervals will contain the true parameter values with >90% probability). For validation, we therefore used a Bayesian simulation scheme where we first draw parameters from their prior distribution, generate data given the drawn parameters, and then fit the Bayesian model with the same prior distributions; by construction, this scheme should match the nominal coverage if the model fits are correct under their own assumptions (Cook et al., 2006). We sampled 100 sets of the parameters from the same prior distribution that was used in the fitting process; for each parameter set, we simulated one realization of 15 time steps (10 for fitting and 5 to compare to forecasts). All model variants were used to fit each realization (Table 1 and 2 in the appendix give more detail about parameters and priors). We added two convergence criteria to assess convergence for the main parameters (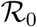, *P*_eff_, *P*_rep_): we required a value of the Gelman and Rubin statistic 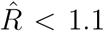 and an effective sample size (ESS) greater than 400 for each replication. For each replication we sample four chains starting with 4000 iterations; we repeatedly double the number of iterations (with a upper threshold of one million iterations) until the convergence criteria are met. Forecasts were made by simulating incidence 5 time steps forward using parameters sampled from the fitted posterior distributions.

**Table 1:**
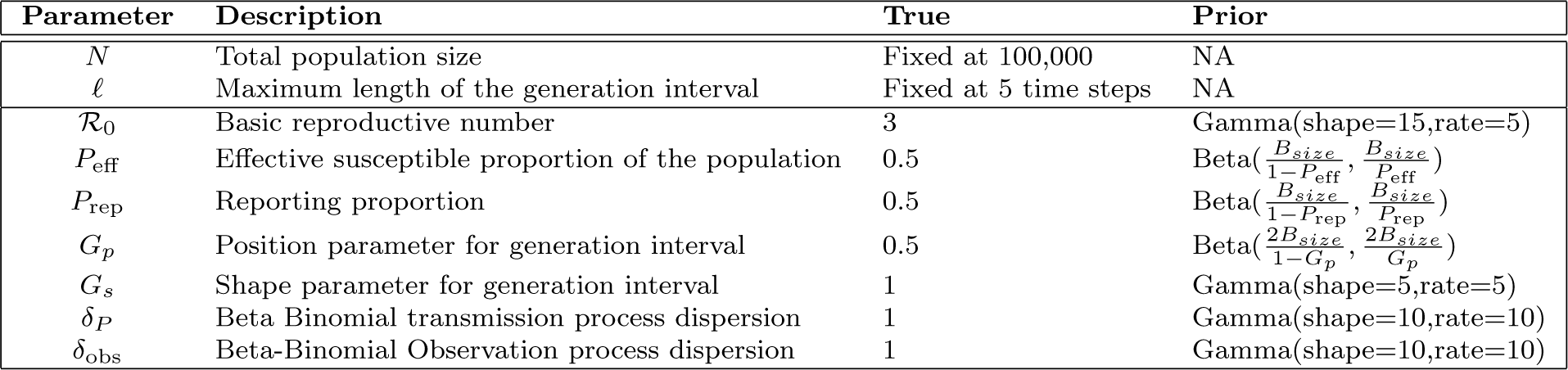
Simulation model parameters

**Table 2:** Fitting model parameters

| Parameter | Description | True | Prior |
| --- | --- | --- | --- |
| $N$ | Total population size | Fixed at 100,000 | NA |
| $\ell$ | Maximum length of the generation interval | Fixed at 5 time steps | NA |
| $B_{\text{size}}$ | Beta prior size factor | Fixed at 1 | NA |
| $\mathcal{R}_0$ | Basic reproductive number | 3 | Gamma(shape=15,rate=5) |
| $P_{\text{eff}}$ | Effective susceptible proportion of the population | 0.5 | Beta($\frac{B_{\text{size}}}{1-P_{\text{eff}}}$, $\frac{B_{\text{size}}}{P_{\text{eff}}}$) |
| $P_{\text{rep}}$ | Reporting proportion | 0.5 | Beta($\frac{B_{\text{size}}}{1-P_{\text{rep}}}$, $\frac{B_{\text{size}}}{P_{\text{rep}}}$) |
| $P_{\text{effrep}}$ | Proportion of effective S to I that are observed | $P_{\text{eff}} \times P_{\text{rep}}$ | Beta($\frac{B_{\text{size}}}{1-P_{\text{effrep}}}$, $\frac{B_{\text{size}}}{P_{\text{effrep}}}$) |
| $\rho$ | Scale splitting factor | 0.5 | Beta($\frac{B_{\text{size}}}{1-\rho}$, $\frac{B_{\text{size}}}{\rho}$) |
| $G_p$ | Position parameter for generation interval | 0.5 | Beta($\frac{2B_{\text{size}}}{1-G_p}$, $\frac{2B_{\text{size}}}{G_p}$) |
| $G_s$ | Shape parameter for generation interval | 1 | Gamma(shape=5,rate=5) |
| $\delta_P$ | Beta Binomial transmission process dispersion | 1 | Gamma(shape=10,rate=10) |
| $\delta_P$ (Neg-Binom) | Negative-Binomial Transmission process dispersion | NA | Uniform(min=0,max=100) |
| $\delta_{\text{obs}}$ | Beta-Binomial Observation process dispersion | 1 | Gamma(shape=10,rate=10) |
| $\delta_{\text{obs}}$ (Neg-Binom) | Negative-Binomial Transmission process dispersion | NA | Uniform(min=0,max=100) |

We evaluated our estimates of (1) total cases predicted over the forecast window (disaggregated forecasts are analyzed in the supplementary material) and (2) key model parameters (including the estimated mean generation interval (MGI: defined 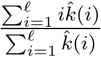)). We used bias, root mean square error (RMSE), and coverage to assess model fit. Bias and RMSE are based on proportional errors, defined as the log ratio of our estimate (taken as the median of the posterior sample) to the known true parameter value from our simulations. Errors were compared on the log scale in order to allow comparison of the accuracy of estimation of different parameters that may be on very different scales. The median is a scale-invariant, robust, summary statistic for the location parameter of a Bayesian posterior distribution Minsker et al. (2014). Thus in order to compare different parameters in a consistent, unitless fashion, the errors were calculated as 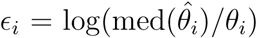. We then calculated bias (median(*ϵ*)) and RMSE 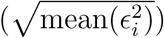.

## 3 Results

The full model (which matches the simulation model) provides generally good forecasts and parameter estimates as assessed by bias (Figure 3) or RMSE (Figure 4), except for estimates of *P*_eff_ using JAGS.

**Figure 3:**
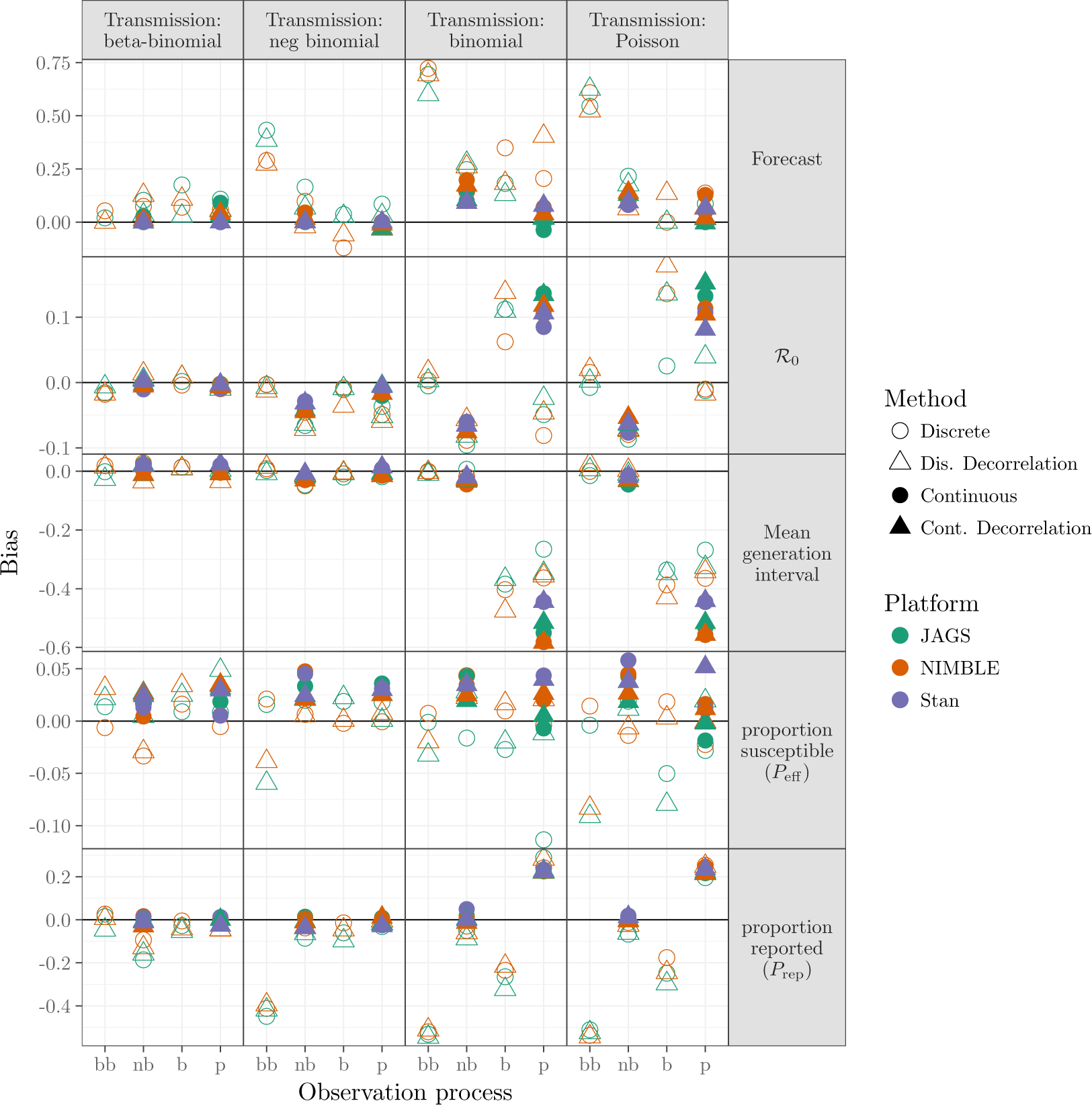
Comparison of bias (based on proportional errors) for forecasts and parameters using models described in Sect. 2.2 across different platforms described in Sect. 2.3.1. Models with overdispersion in the transmission process (BB and NB, leftmost and second-left columns of panels) and models with overdispersion in the observation process (BB and NB, leftmost and second-left x-axis ticks within each panel) have generally low bias. Continuous latent-state models (solid points) are only implemented for negative binomial and Poisson observational processes.

**Figure 4:**
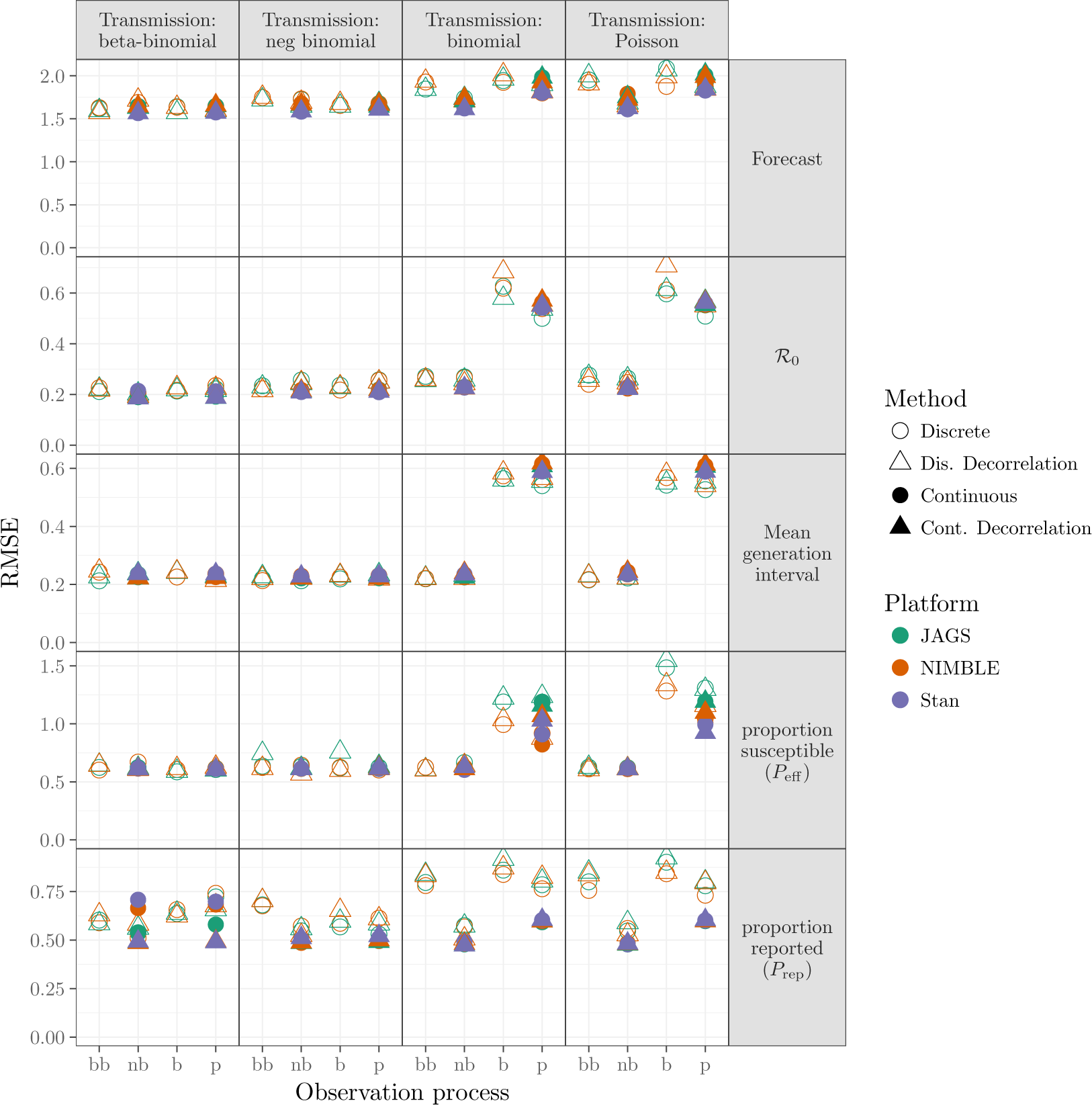
Comparison of RMSE (based on proportional errors) for all fitting model variants. The layout matches that of Figure 3. Patterns across models and platforms are similar to those seen in Figure 3. Short-term forecasts have generally high error, even when bias is low, reflecting inherent uncertainty in the system. The highly correlated parameters *P*_eff_ and *P*_effrep_ also show high error but not high bias.

In general, models with any kind of dispersion in the transmission process, or with negative binomial dispersion in the observation process, did well. The exception is that models that combined negative binomial transmission dispersal with beta binomial observation dispersal produced biased forecasts and estimates of *P*_rep_.

There are no clear differences in the quality of model fit due to multi-scale decorrelation, latent continuous transmission process or platform.

Figure 5 shows the statistical coverage of our estimates. Similar to the results for bias and RMSE (Figure 3 and 4), we find generally good coverage (i.e., close to the nominal value of 0.9) for models with dispersion in the transmission process, except that the negative-binomial transmission process model undercovers across the board (coverage ≈ 0.8 for all observation process models and platforms) for forecasts and *P*_rep_. For models without dispersion in transmission, models with dispersion in the observation process have low coverage (≈ 0.8) for most parameters, while the betabinomial process model has low coverage (≈ 0.>4) for *P*_rep_ and models without any dispersion have uniformly low coverage.

**Figure 5:**
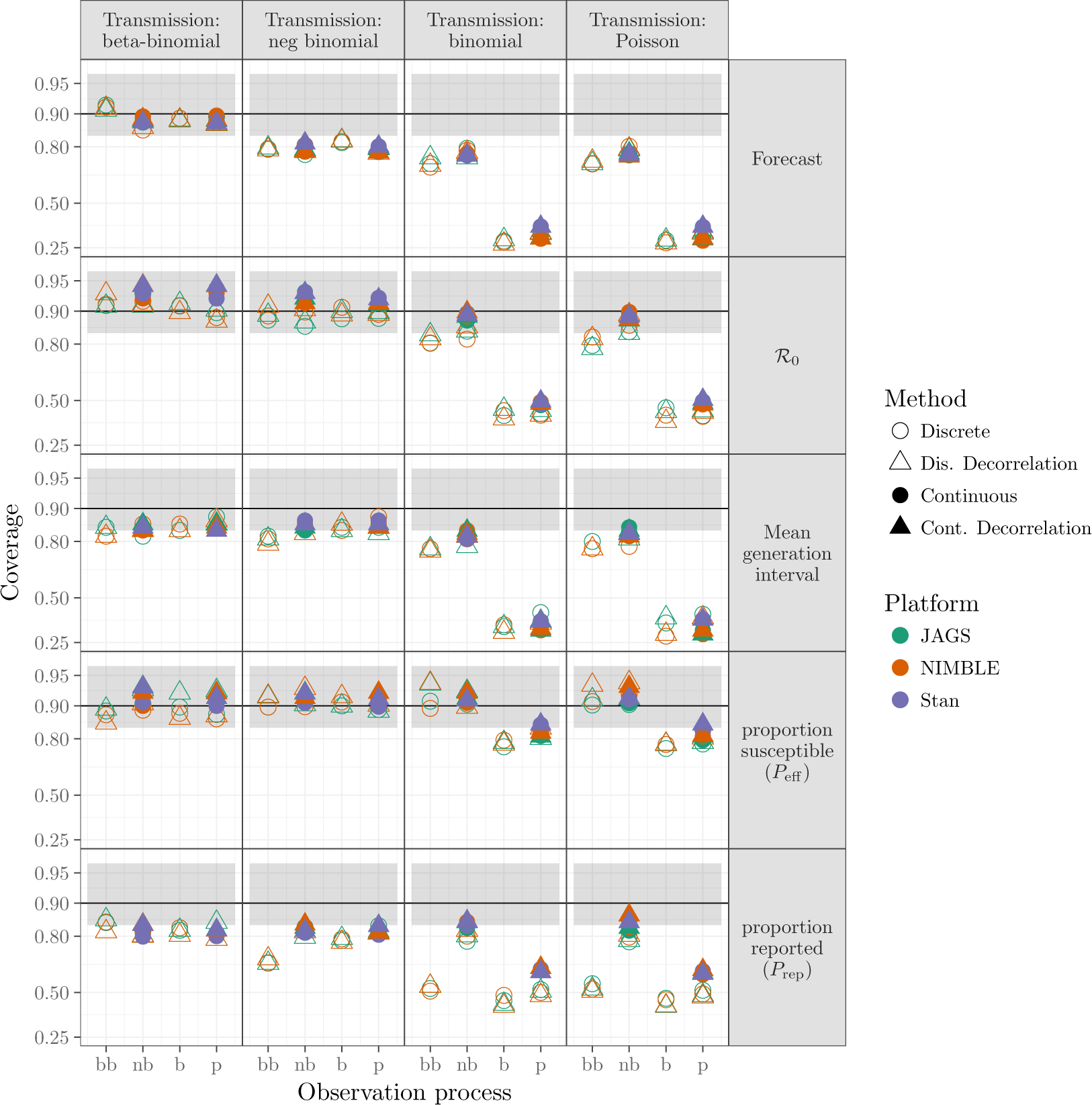
Comparison of coverage probability for forecast and parameters. Models with overdispersion in the transmission process (BB and NB, leftmost and second-left columns of panels) and models with overdispersion in the observation process (BB and NB, leftmost and second-left x-axis ticks within each panel) have coverage near the nominal value of 0.9 for all parameters and model variants. The black line shows the nominal coverage, and the grey ribbon the 95% binomial confidence interval based on 100 simulated fits. Vertical axis is plotted on a logit scale.

There are substantial efficiency differences between transmission-process approaches (continuous vs. discrete), as measured by time per effective sample size, shown in Figure 6. For a given platform, models using continuous latent variables are generally more efficient than discrete latent processes. Comparing models with continuous latent variables between platforms (Figure 5, second and fourth column of every panel), Stan (using HMC) is sightly more efficient for majority of the parameters, followed by NIMBLE and JAGS. Furthermore, continuous latent-variable models (especially using HMC in STAN) use fewer iterations (when meeting all convergence criterion described in section 2.3.2) than discrete latent-variable models.

**Figure 6:**
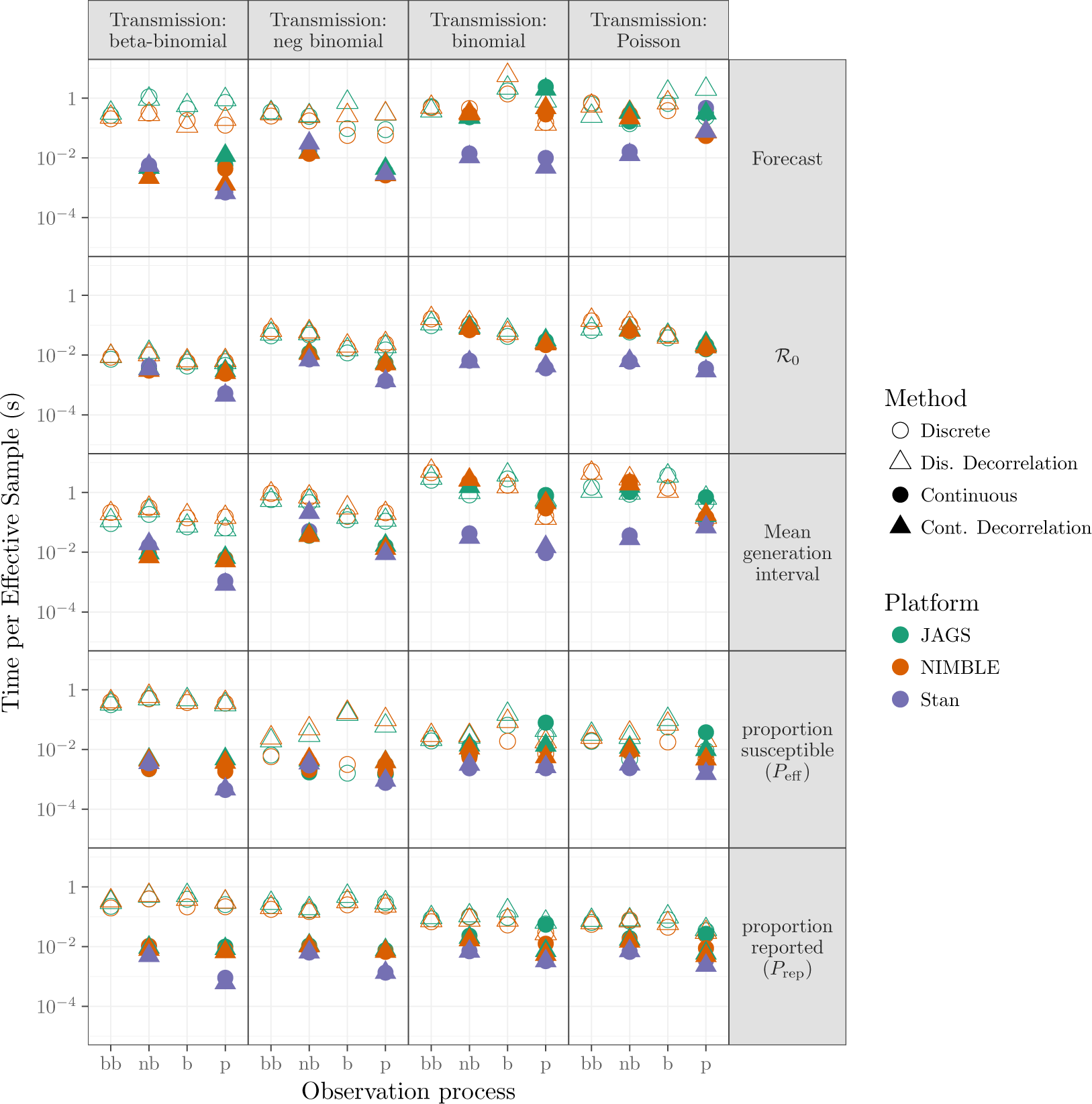
Comparison of efficiency for all fitting model variants: layout of models and platforms as in Figure 3.

## 4 Discussion

We have fitted models varying in complexity to simulated epidemic data with multiple sources of heterogeneity, using several different platforms. Using models that include some form of overdispersion is necessary for robust fits, but models that include overdispersion only in the transmission process can work as well as or better than the full model. Including overdispersion only in the observation process (if implemented as a negative binomial distribution) also provides relatively robust fits to these data. Simplifying the models by using continuous rather than discrete latent variables increased efficiency with little effect on the quality of the fits.

### 4.1 Ceilings

The effects of using distributions with ceilings (i.e. binomial and beta-binomial distributions) instead of their less realistic counterparts without ceilings (Poisson and negative binomial) was relatively small. In our framework, ceilings only apply in models with discrete latent variables; the primary effect of such ceilings is to reduce variance as probabilities (of infection or of sampling) become large. (Reporting-process models without ceilings also allow for false positives or over-reporting, which may be important in some contexts.)

### 4.2 Overdispersion

Accounting for overdispersion had more impact on our fits than the presence or absence of ceilings. In particular, models with no overdispersion in either process lacked flexibility and tended to be over-confident (that is, they showed low coverage). However, models that account for overdispersion in only one process (either transmission or observation) tended to be reliable for estimating parameters such as 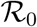, mean generation interval, and short-term forecasts, particularly when overdispersion was implemented through negative binomial (a less constrained distribution than the beta binomial). However, parameters that are closely tied to the details of a particular model structure (such as the overdispersion parameters for the observation and transmission processes) must change when the overdispersion model changes, in order to compensate for missing sources of variability.

Several authors (e.g., (King et al., 2015; Taylor et al., 2016)) suggest that accounting for process as well as observation error in estimates of 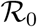 and in forecasts is necessary in order to avoid over-confident estimates. Our exploration does not include any cases where process error is completely absent—even our “dispersion-free” processes incorporate sampling error in the process. However, we find that neglecting overdispersion can still lead to over-confident and unreliable estimates.

### 4.3 Reporting

In classic infectious disease models, reducing reporting rate and reducing the total effective population size have similar effects: reducing the observed size of the epidemic. While we want to make as few assumptions as possible about unobservable aspects of the epidemic, underreporting is of huge practical importance. Additionally, modeling observation error explicitly is required for reliable estimates of uncertainty (King et al., 2015). If reporting error is modeled with a ceiling, then underreporting is a necessary component of reporting error (i.e., reporting is always biased downward as well as noisy). Allowing overdispersion decouples the variance from the mean of the reporting process (i.e. the extra overdispersion parameter means that the variance is not determined by the mean).

Because reporting rate and effective population size play similar roles in epidemic dynamics, incorporating them both in a model may make their parameter estimates strongly correlated and hence difficult to identify: we may be very uncertain whether low observed epidemic incidence is driven by a small effective population size or a low reporting rate. We have addressed convergence problems arising from this issue by reparameterizing the model (Section 2.2.2). From a conceptual point of view, joint unidentifiability is not necessarily a serious problem, as long as the quantities we are most interested (such as 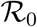) are identifiable. In practice, however, weak identifiability can cause hard-to-detect convergence problems; known-parameter simulations like those implemented here are useful for validation in such cases.

### 4.4 Extensions and alternative approaches

Our analysis covers classical MC (i.e. conditional updating of parameters via conjugate, slice, and Metropolis-Hastings samplers) and HMC approaches. Even within this scope there is additional room for analysis, both in terms of exploring important heterogeneities that we have neglected here (such as spatial, age and social structure), and in improving sampling techniques (e.g. by adjusting the choice of samplers in JAGS or NIMBLE or by redundant parameterization Gelman et al. (2014)).

More broadly, a plethora of other model-fitting tools is available to researchers, from deterministic optimization tools based on the Laplace approximation Illian et al. (2012); Kristensen et al. (2016) to sequential techniques such as iterated filtering and particle MC Del Moral et al. (2012); He et al. (2009); Yang et al. (2014). Ionides et al. (2006). These techniques can in principle be combined flexibly with the methods we explore here, e.g. using HMC to sample top-level parameters while implementing a sequential MC technique for the latent states. It will be interesting to see how the single-technique methods here compete with hybrid approaches, and how flexible toolboxes such as NIMBLE will fare against more focused platforms like Stan.

### 4.5 Prior distributions

This paper focuses on evaluating Bayesian methods for fitting and forecasting epidemics. For the purposes of evaluation we use parameter distributions for simulation that exactly match our Bayesian priors. We are assuming that researchers have a reasonable method of choosing appropriate Bayesian priors; in real applications this will be an important challenge.

## 5 Conclusion

We have presented a comparison of simple MCMC approaches to fit epidemic data. We learned two things about fitting epidemic data. First, modeling different processes with dispersion (BB and NB) is a naive but effective way to add uncertainty in the model; models that neglect such uncertainty are likely to be over-confident and less accurate at forecasting. Second, approximating discrete latent state process with continuous processes can aid efficiency without losing robustness of fit. This allows more efficient fitting in the classic framework (e.g., JAGS and NIMBLE), and also allows us to use the more advanced HMC technique (which we implemented via Stan).

## 6 Acknowledgments

We would like to thank the Ebola challenge organizers for organizing the Ebola model challenge that sparked our interest in this project, and Fred Adler and Michael Betancourt for thoughtful comments. McMaster University, NSERC Discovery grants and a CIHR Ebola grant provided funding.

## Supplementary Material

In the main text, we present the bias, RMSE, coverage and efficiency plots for aggregated forecast, 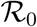, MGI, *P*_eff_, and *P*_rep_. Here, we present plots showing the other parameters (shape *G_S_* and position *G_P_* of the transmission kernel and process and observation overdispersion parameters *δ_P_* and *δ*_obs_) and disaggregated forecasts (five forecast steps) that are excluded in the main text. We also add some representative plots of the simulated cases and forecast.

**Figure S1:**
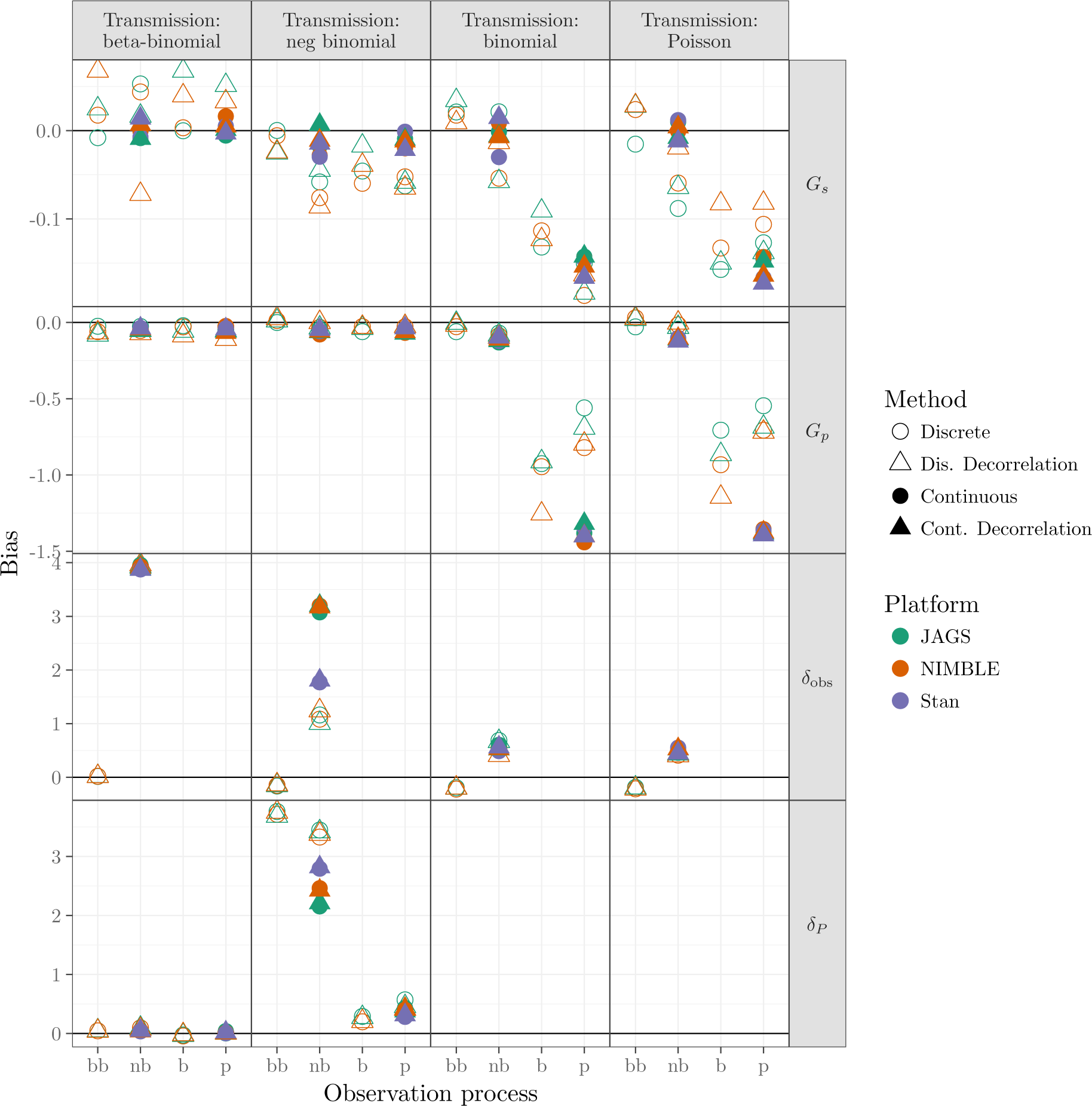
Comparison of bias for *G_S_* (transmission shape), *G_P_* (transmission position), *δ*_obs_ (observation overdispersion), and *δ_P_* (process overdispersion: more detail given in Sect. 2.2) across different platforms (described in Sect. 2.3.1). Overdispersion parameter *δ_P_* is only applicable in models with dispersion in the transmission process (first and second left column panel) and overdispersion parameter *δ*_obs_ is only applicable in models with dispersion in the observation process (first and second column within each column panel).

**Figure S2:**
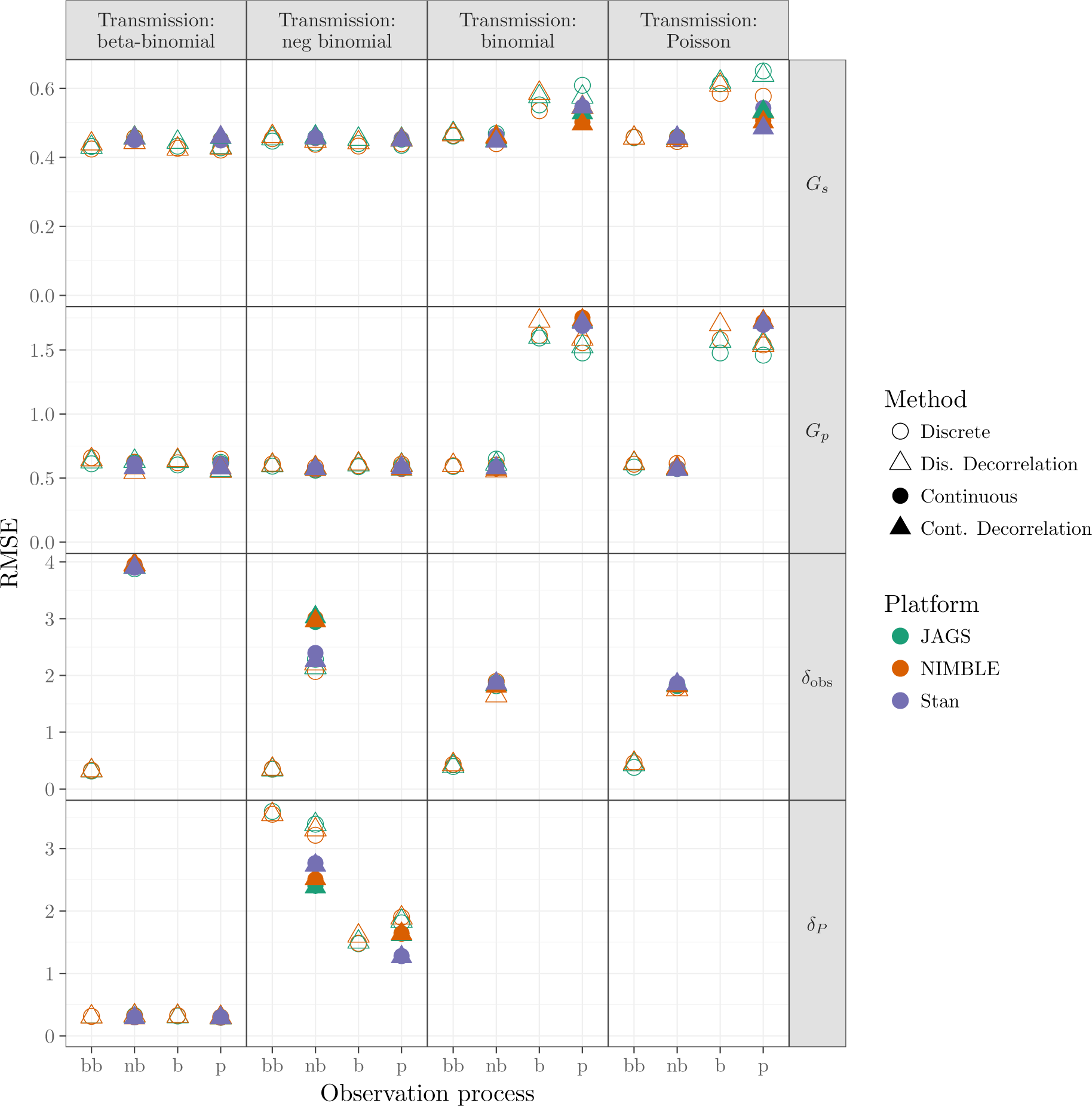
Comparison of RMSE for *G_S_*, *G_P_*, *δ*_obs_, and *δ_P_*. See Figure 4 in main text and Figure S1 in appendix for details.

**Figure S3:**
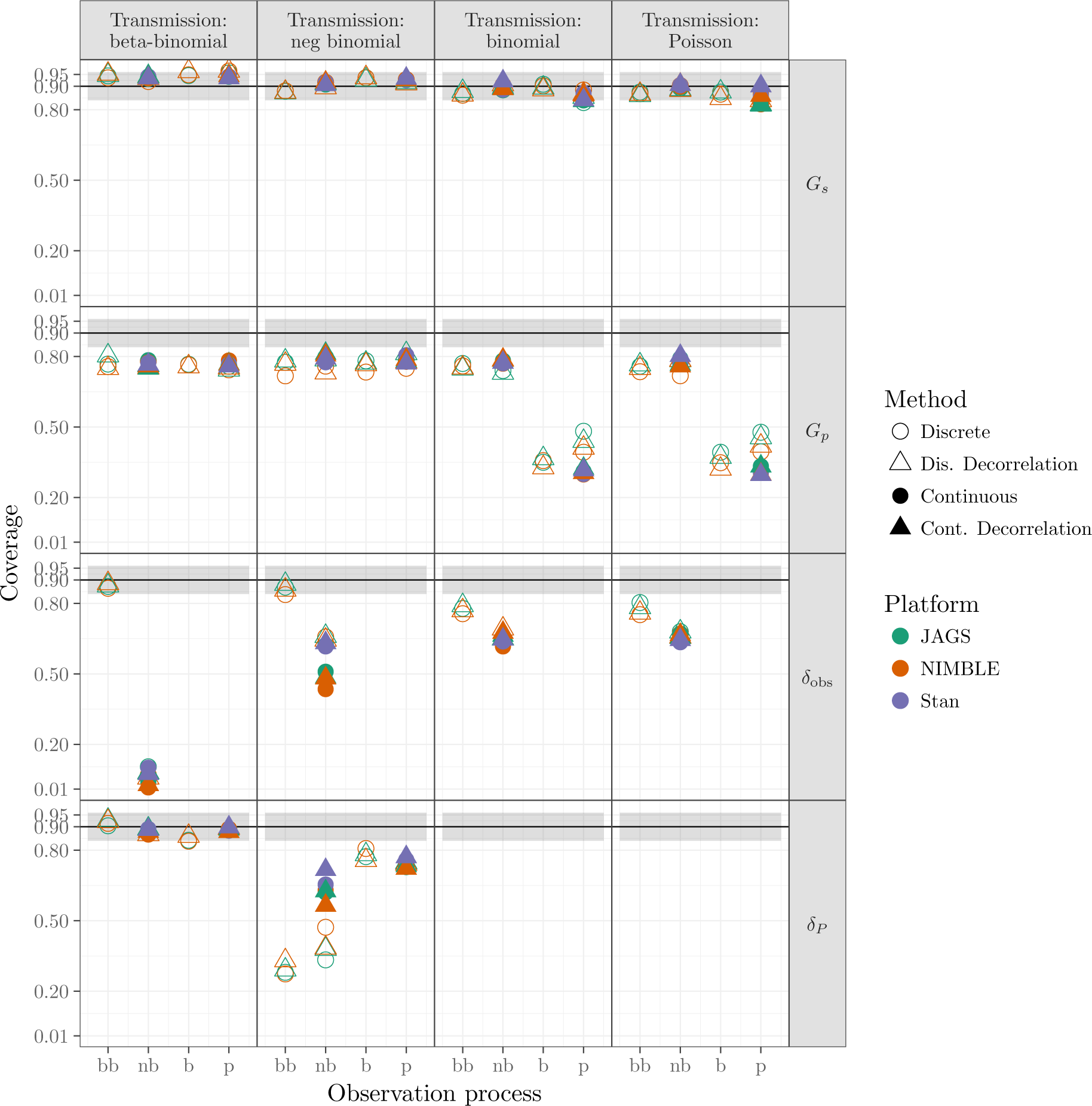
Comparison of coverage for *G_S_*, *G_P_*, *δ*_obs_, and *δ_P_*. See Figure 5 in main text and Figure S1 in appendix for details.

**Figure S4:**
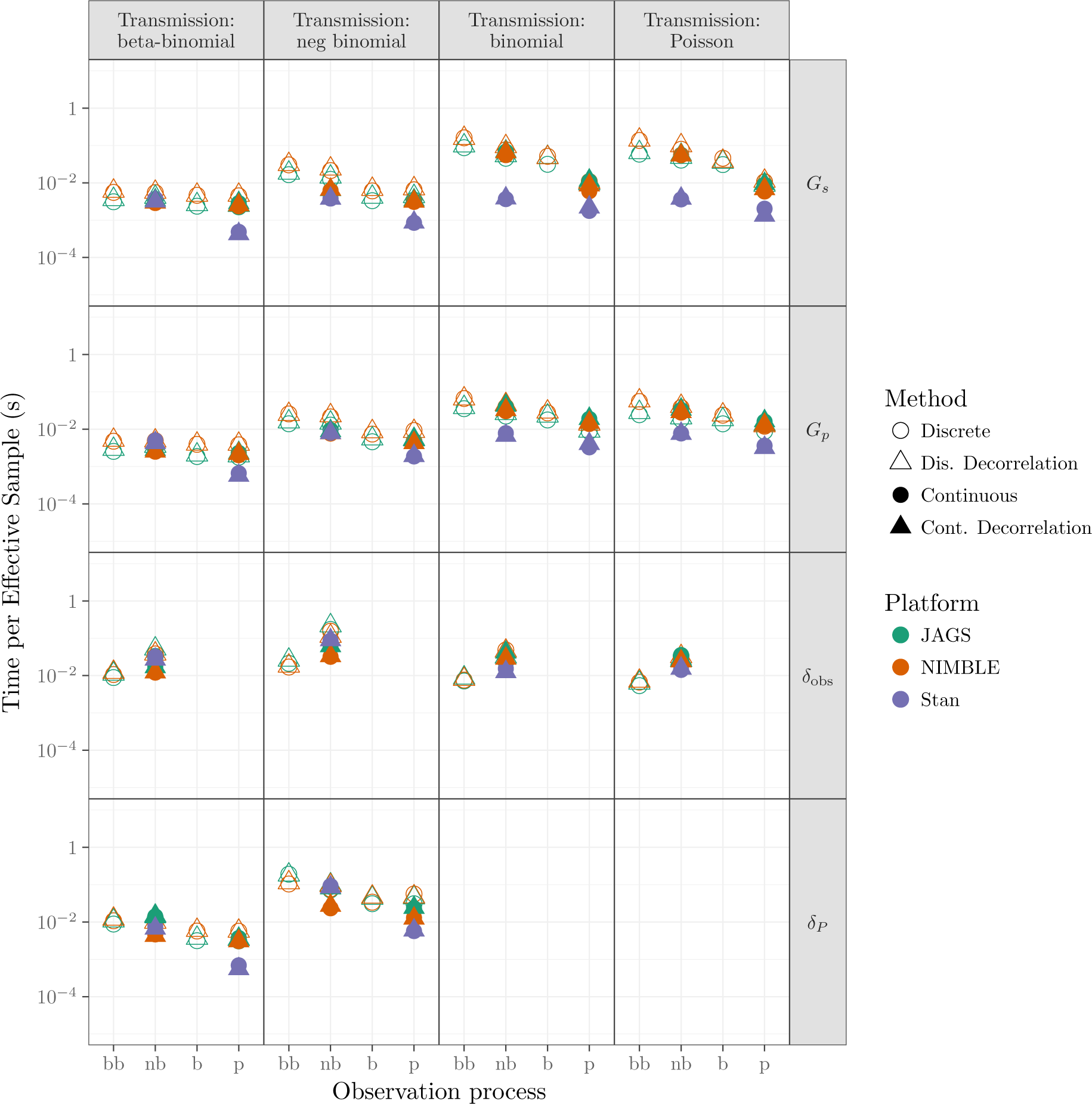
Comparison of coverage for *G_S_*, *G_P_*, *δ*_obs_, and *δ_P_*. See Figure 6 in main text and Figure S1 in appendix for details.

**Figure S5:**
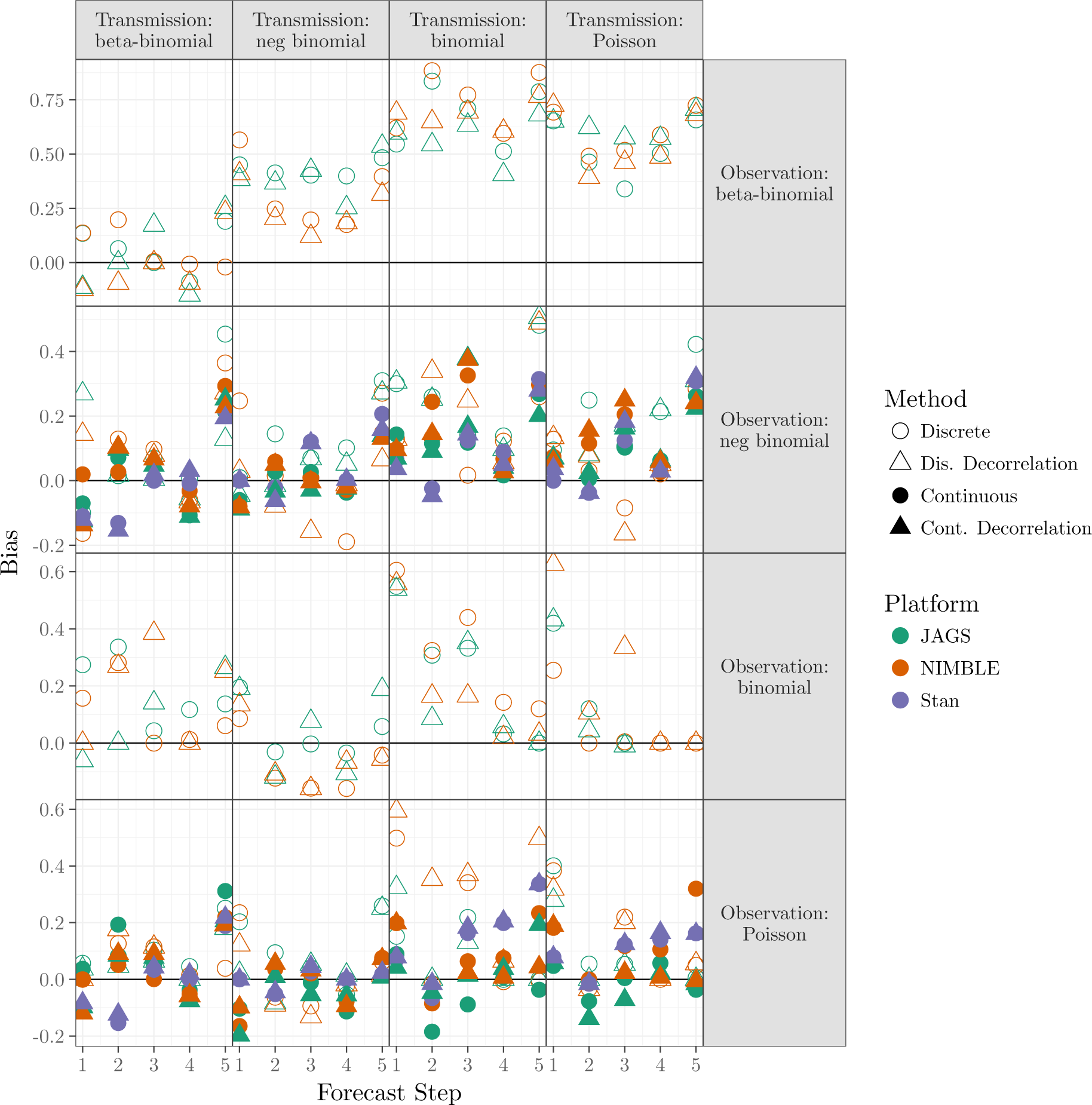
Comparison of bias for five forecast steps (described in Sect. 2.2) across different platforms (described in Sect. 2.3.1).

**Figure S6:**
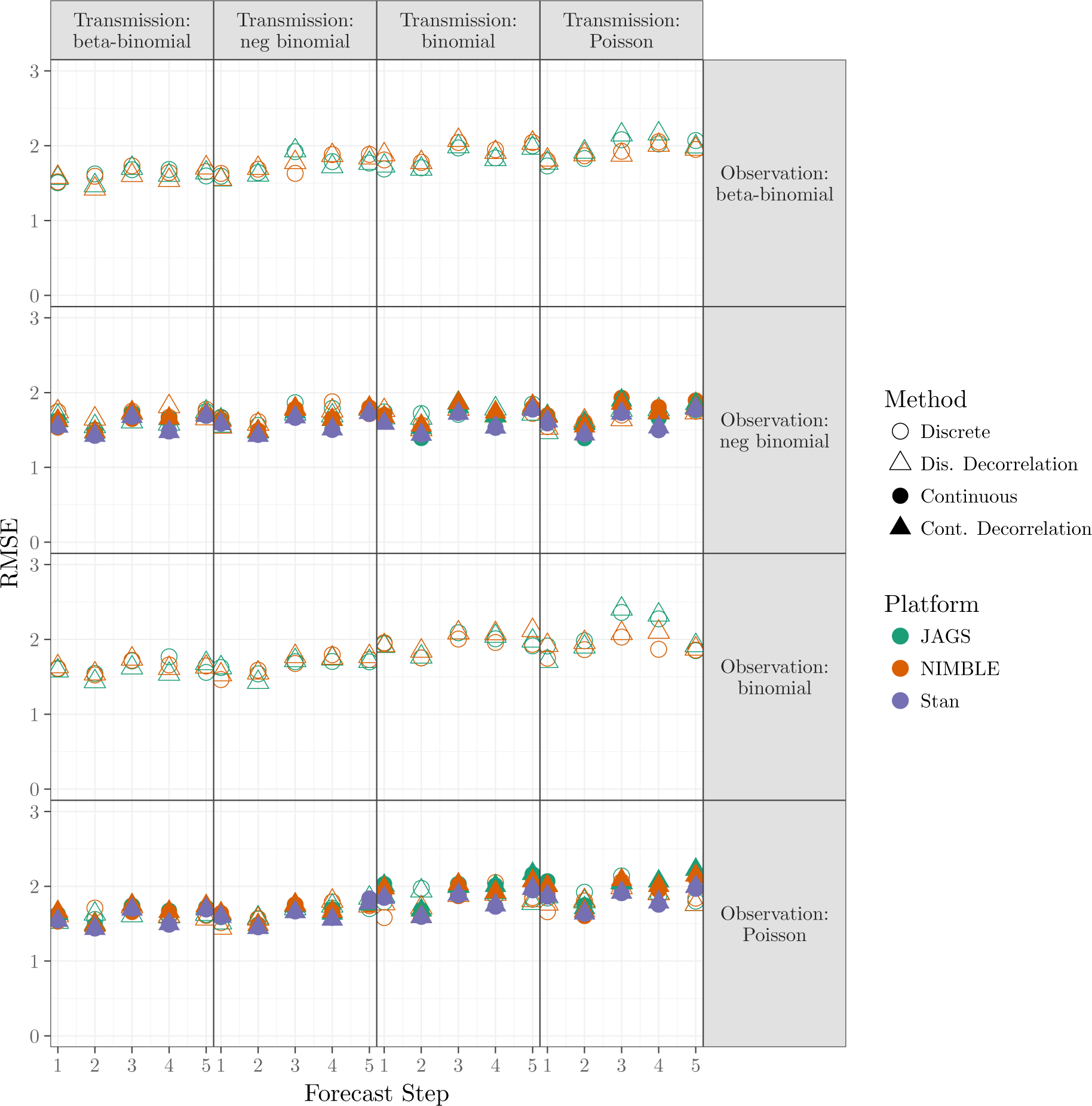
Comparison of RMSE for five forecast steps described in Sect. 2.2 across different platforms described in Sect. 2.3.1. See Figure 4 in the main text for details.

**Figure S7:**
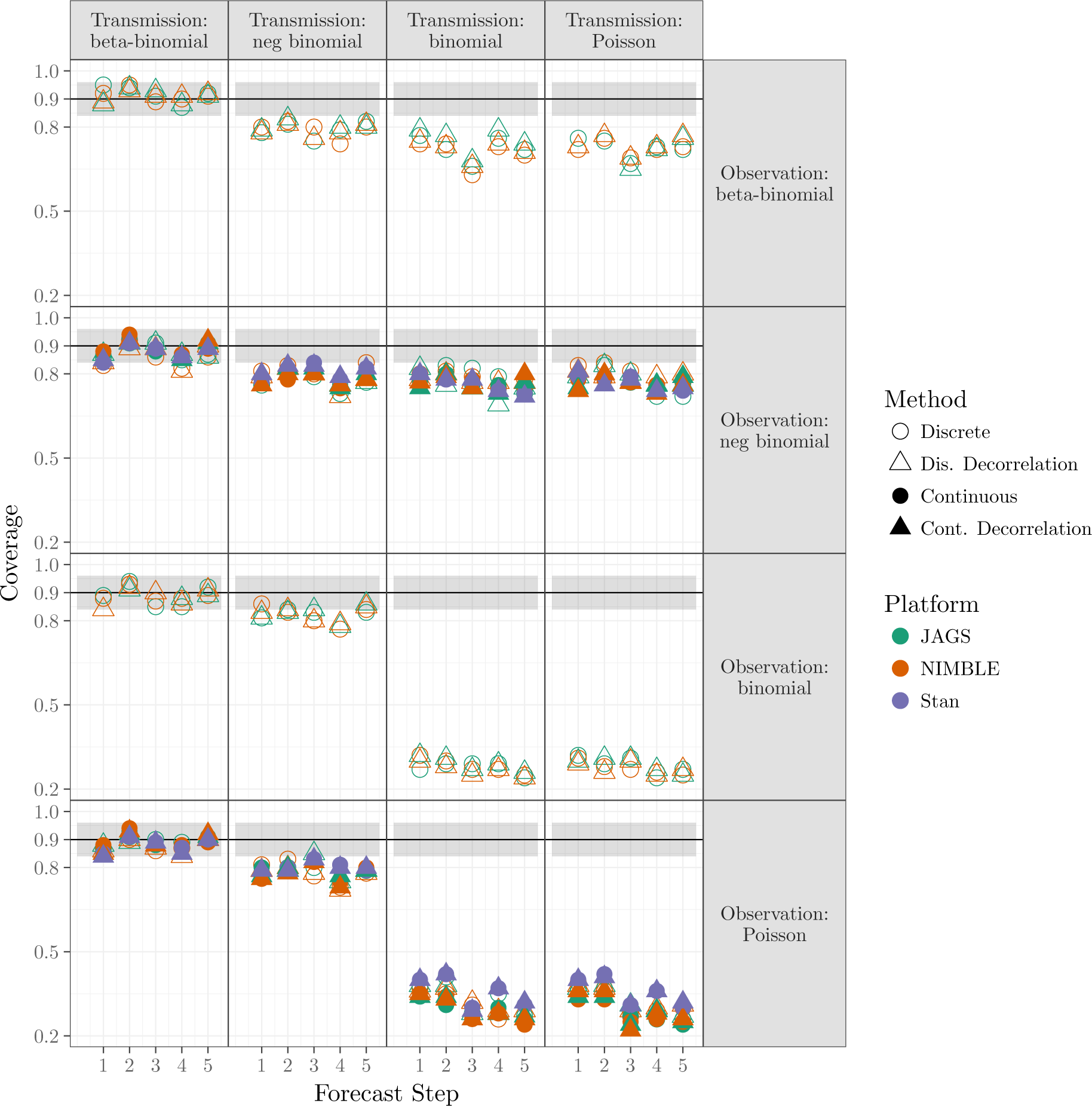
Comparison of coverage for five forecast steps described in Sect. 2.2 across different platforms described in Sect. 2.3.1. See Figure 5 in the main text for details.

**Figure S8:**
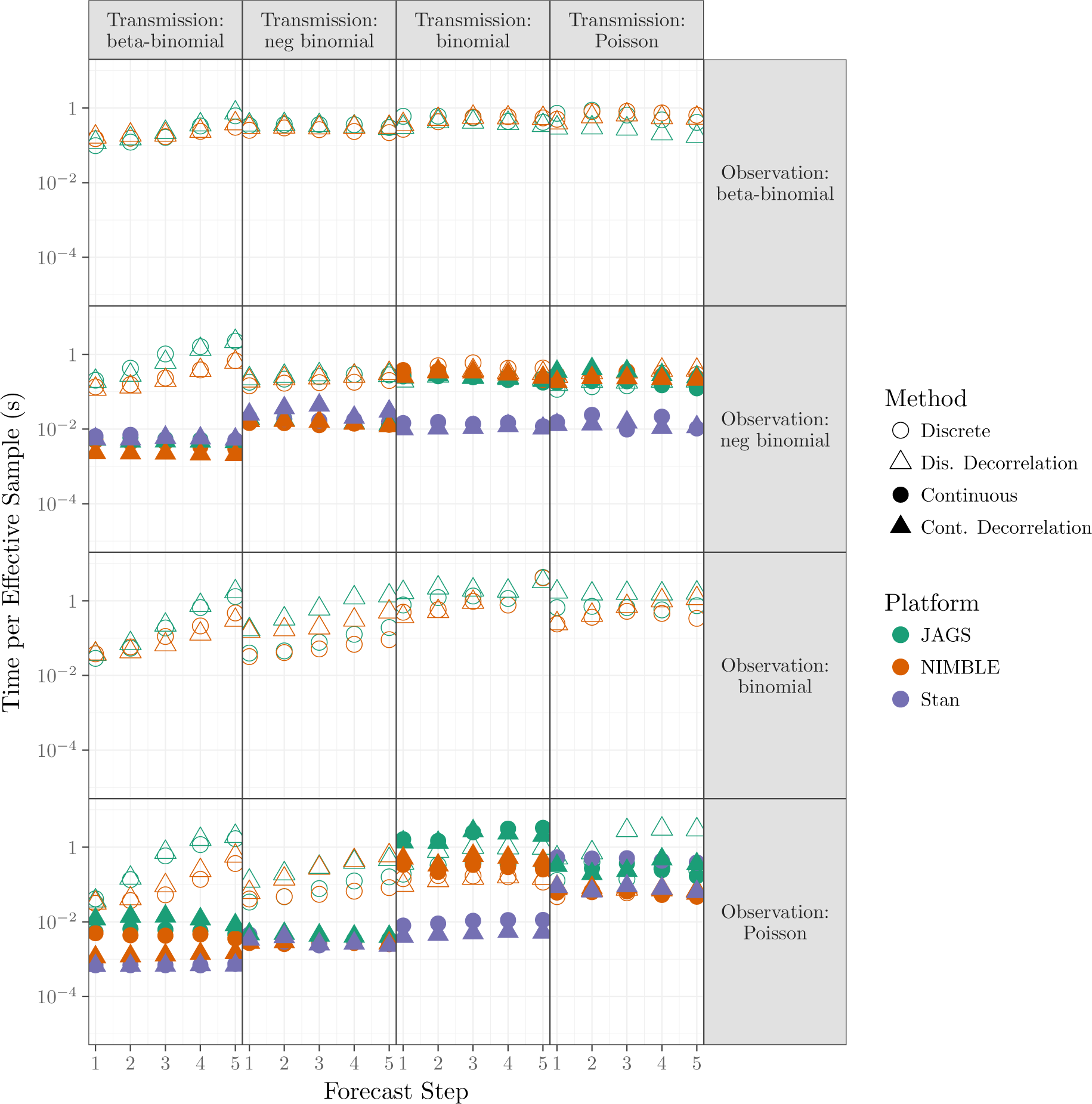
Comparison of sampling efficiency for five forecast steps described in Sect. 2.2 across different platforms described in Sect. 2.3.1. See Figure 6 in the main text for details.

**Figure S9:**
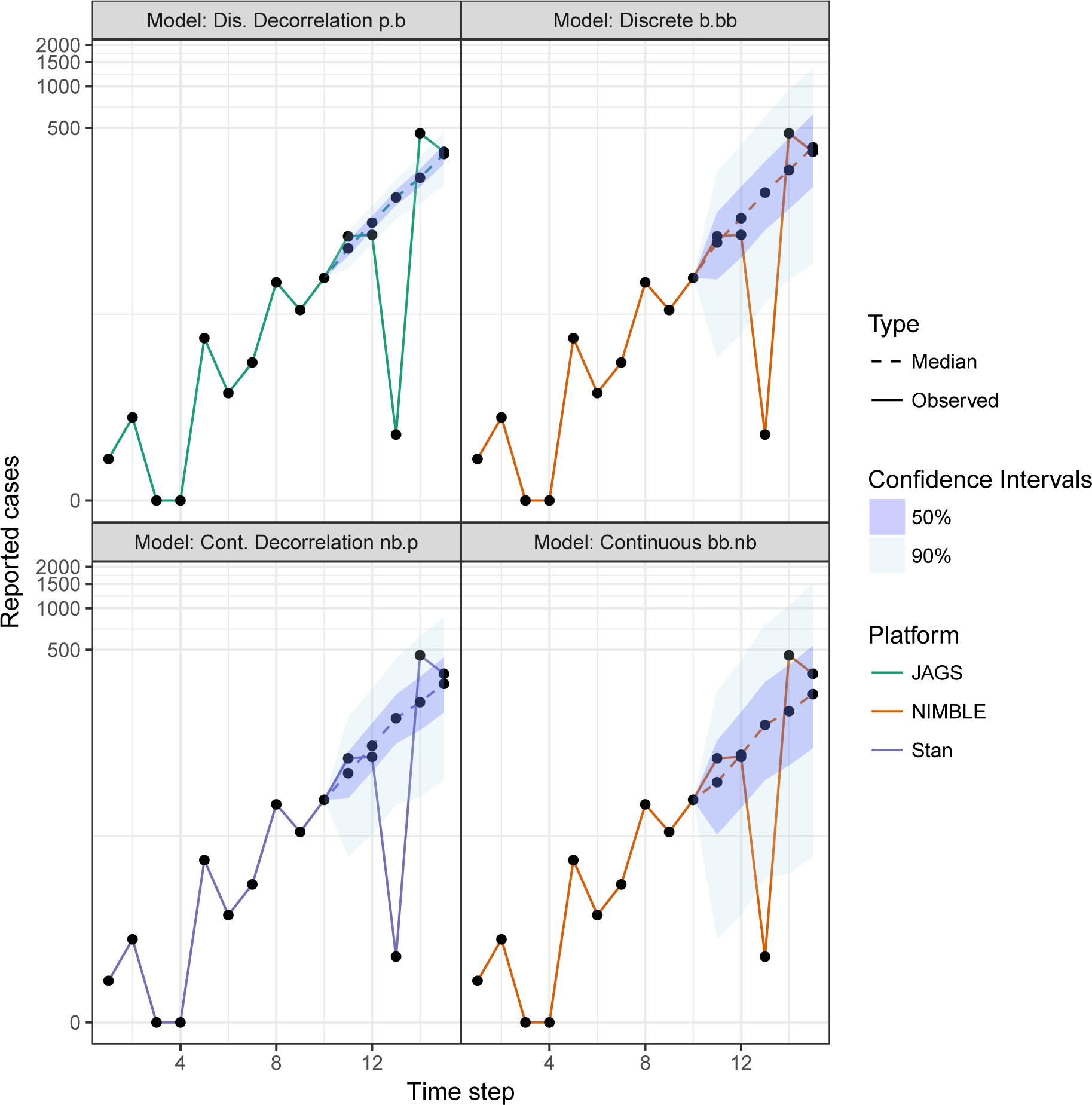
Comparison of forecast using combinations of transmission process, observation process, decorrelation, latent state variables, and platforms described in Sect 2.2 and 2.3.1. Moving from the top to bottom row adds overdispersion in the transmission process (binomial (b) and Poisson (p) to negative-binomial (nb) and beta-binomial (bb)). Moving from left to right adds overdispersion in the observations. Solid line shows the simulated observed cases (15 time steps); dashed line shows the median of the posterior forecast sample with 50% (dark ribbon) and 90% (light ribbon) confidence intervals (last 5 time steps).

**Figure S10:**
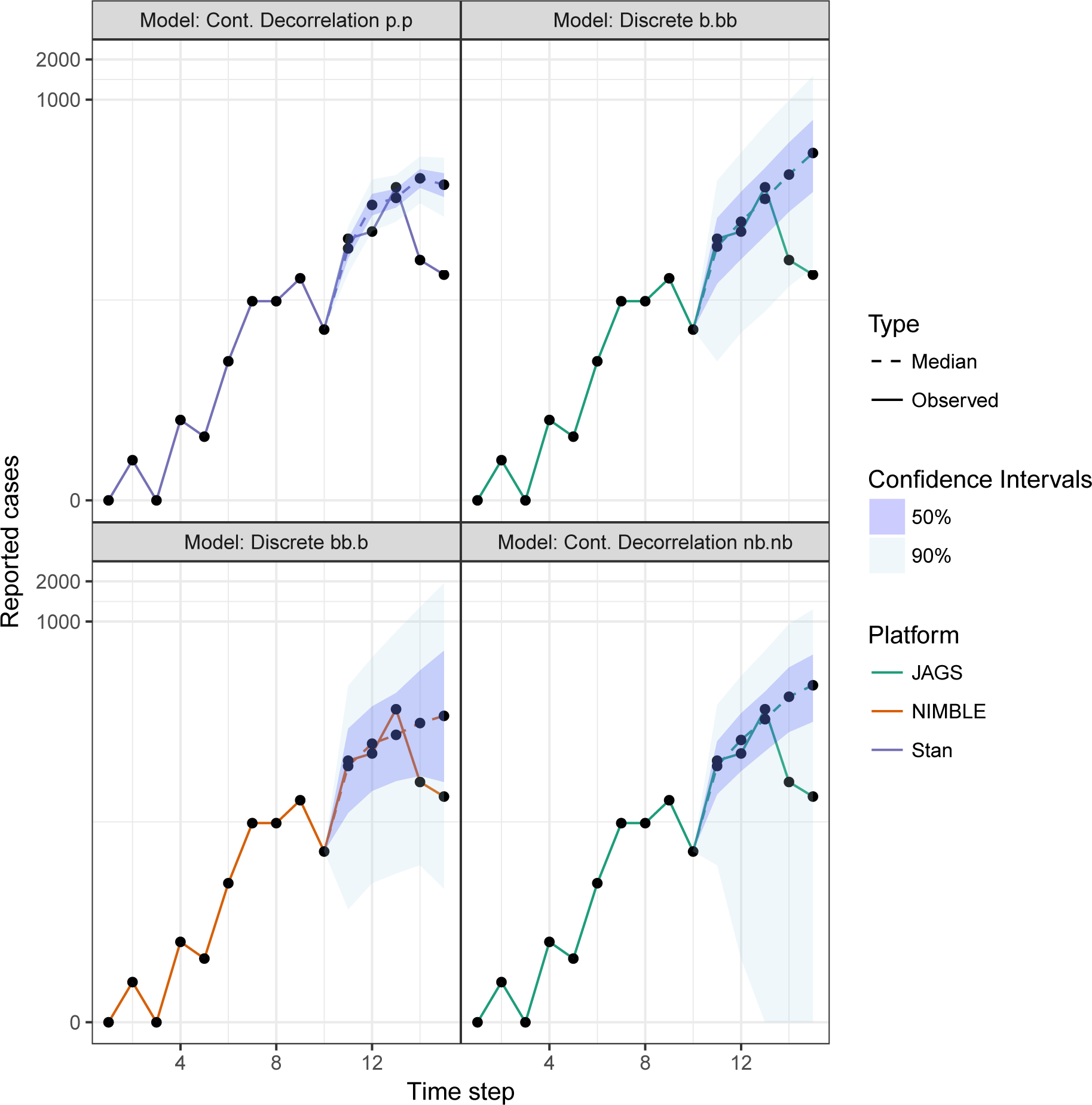
Comparison of forecast using a different set of parameters. See Figure S9 for details.

**Figure S11:**
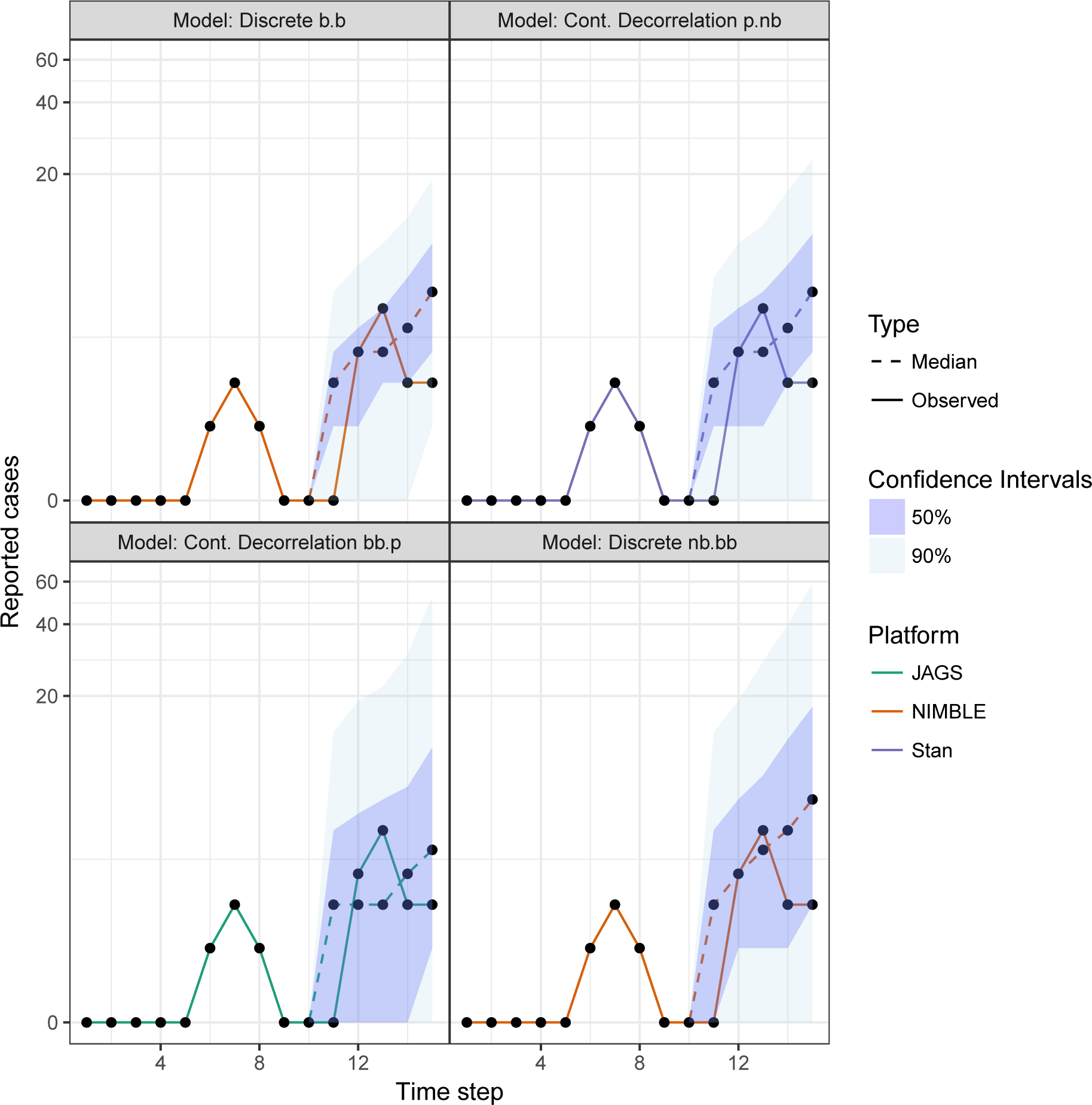
Comparison of forecast of low observed cases. See Figure S9 for details.

